# Electromotility can be disassociated from gating charge movement in outer hair cells of conditional alpha2 spectrin knockout mice

**DOI:** 10.1101/2025.02.25.640135

**Authors:** Jun-Ping Bai, Micheal Stankewitch, Jie Yang, Winston Tan, Zhongyuan Zuo, Qiang Song, Saaim Khan, Jon Morrow, Joseph Santos-Sacchi, Dhasakumar S. Navaratnam

## Abstract

Electromotility in mammalian outer hair cells (OHC) is the mechanism underlying cochlear amplification. It is brought about by the piezoelectric-like property of the membrane protein prestin (Slc26a5) that lies in the OHCs lateral plasma membrane. Prestin connects to an underlying cytoskeletal network of circumferential actin filaments that bridge longitudinal spectrin filaments. This network, in turn, lies between the plasma membrane and a closely apposed ER-like tubular array of subsurface cisternae (SSC). Two previous papers examining spectrin knockouts in embryonic hair cells were confined to analyzing the effects on the apical cuticular plate and overlying stereocilia. In this paper, we examine the effects of conditional knockouts of alpha2 spectrin in postnatal OHCs. We find a significant auditory phenotype likely due to the novel disassociation of prestins gating charge movement from OHC electromotility. In addition, OHCs show enlargement in their SSC and plasma membrane-SSC space with preserved cuticular plates and overlying stereocilia, which contrasts with the findings in embryonic knockouts.

## INTRODUCTION

Outer hair cells (OHCs) in the mammalian cochlea are responsible for cochlea amplification that is responsible for mammalian hearing sensitivity (*1*). Cochlea amplification is brought about by the OHCs unusual voltage dependent electromechanical coupling known as electromotility (*1, 2*). It is widely believed that electromotility is a membrane-based phenomenon (*3–5*), and the identity of prestin, a member of the Slc26 family of membrane-based anion transporters, as responsible for electromotility cemented this idea (*6*). Prestin is found in high density in the lateral plasma membrane of OHCs (*6*) and replicates all of the electrophysiological characteristics of the OHCs membrane motor (*6–8*). These include the many features of its voltage dependent gating charge movement measured as a non-linear increase in membrane capacitance (NLC) (*6–8*).

OHCs are unusual in that they contain an ER like network of cisternal membranes beneath the lateral plasma membrane (*9–11*). These subsurface cisternae are interconnected with the overlying lateral membrane by a dense cytoskeletal network of longitudinal spectrin and circumferential actin filaments that in turn are linked to the plasma membrane by electron dense pilar like structures (*12–15*). The identity of the specific spectrin pairings or the pilar like structures and how these cytoskeletal elements are involved in electromotility remains indeterminate.

The cytoskeleton plays a critical role in electromotility in two biophysical models for electromotility including the area motor model (*16–18*) and the membrane bending model (*19, 20*). In the area motor model, expansion of the intramembrane motor induces differing stressors in the circumferential and longitudinal direction through the cytoskeleton . Since the axial stiffness is less than the circumferential stiffness, force is preferentially directed along the longitudinal axis through extensible spectrin filaments (*16, 17, 18 , 21*)}. In the membrane bending model, force generation depends on resistance of the cytoskeleton to membrane bending (*19, 20, 22*). Reducing membrane curvature between successive pilars arrayed in the longitudinal axis leads to increased spacing between pillars and elongation of the extensible spectrin filaments that are arranged longitudinally. Both these models involve spectrin filaments playing a critical role in electromotility. Consistent with this framing, diamide, an inhibitor of spectrin filament formation, results in reduced force generation (*21*).

Spectrin was identified as likely to be the longitudinal filaments in the lateral wall by its size (3-5nm) (*13, 23*), and then confirmed by immunolabelling (*12, 24, 25*). Non erythroid alpha spectrin was identified in the lateral wall of OHCs using immunoelectron microscopy (*25*). Recently, two papers discussed effects of knocking out αΙΙ spectrin and βII spectrin in hair cells Key findings in these papers that used two early hair cell transcription factors (Atoh1 starting at E 14.5 and Gfi1 starting at E 16.5) to drive Cre recombinase to induce hair cell specific knock outs were largely descriptive of the effects on the cuticular plates where spectrin is densely expressed and resultant distortions to the overlying stereociliary morphology (*26, 27*). In this paper, we use an inducible prestin-Cre (*28*) to delete αΙΙ spectrin (*29*) in postnatal OHCs and explore the effects of the deletion on electromotility.

## RESULTS

### αΙΙ spectrin is found in the lateral wall of OHCs

We used knockout validated antibodies against αΙΙ spectrin to determine the localization of αΙΙ spectrin in OHCs. Fluorescence immunolocalization of fixed mouse cochlea demonstrated that αΙΙ spectrin was distributed along the lateral wall in proximity to prestin (Figures 1). These data confirm prior findings in the mouse and guinea pig including EM localization of the protein (*12, 24, 25*). In addition to localization along the lateral wall, we also confirm dense expression of αΙΙ spectrin in the cuticular plate of inner and outer hair cells. These latter findings were confirmed in the recent papers on embryonic knockouts of spectrin in mice (*26, 27*).

**Figure 1.**
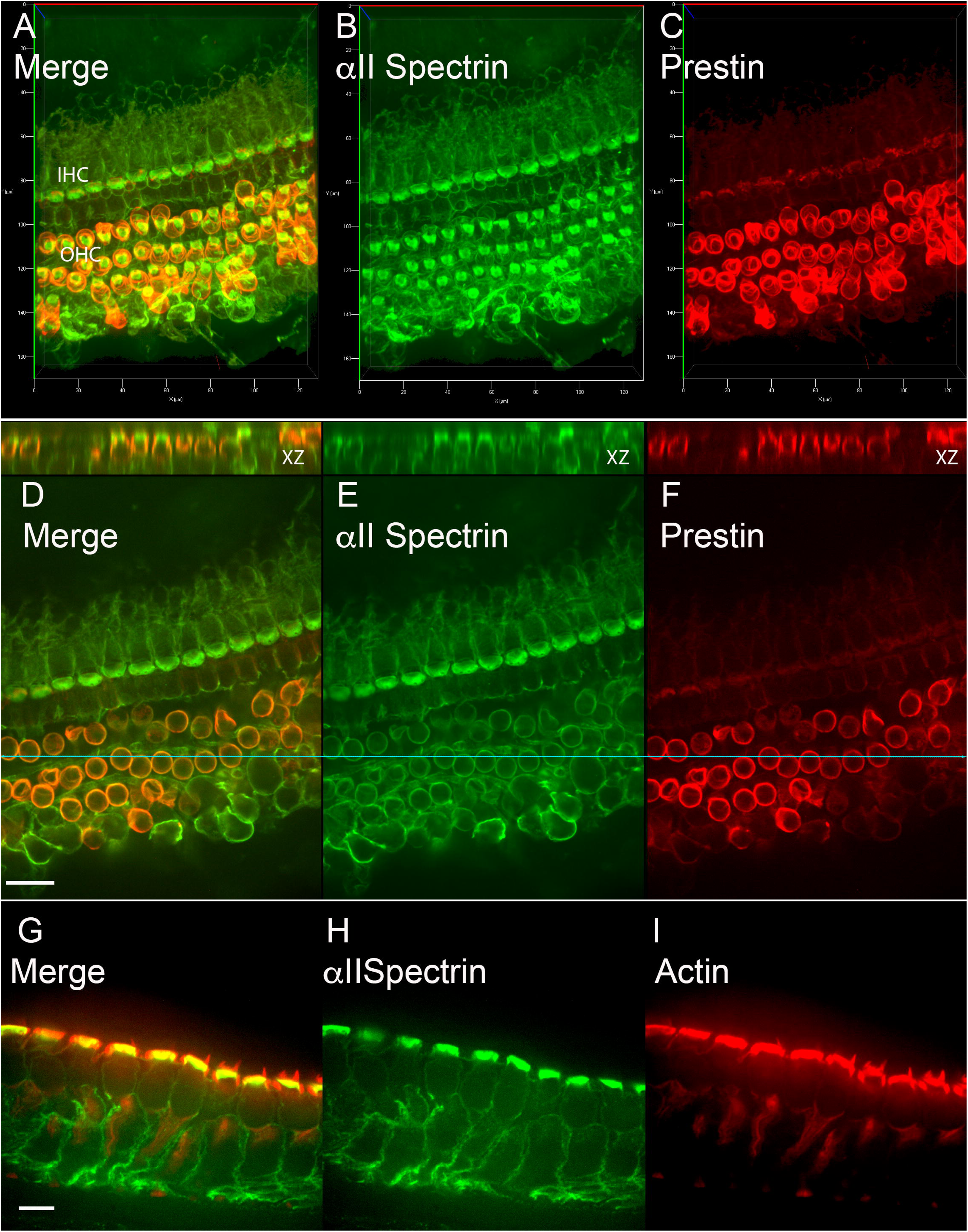
αII spectrin is expressed in the lateral wall of OHCs alongside prestin and b actin. A-C: 3D projection of a z stack from confocal images of IHC and OHC of WT cochlea immunolabeled with antibodies to αII spectrin and prestin (A merged, B a spectrin and C prestin). Note the equal αII spectrin in the cuticular plates of IHC and OHC.D-F. Shown are images of the identical cochlea, now with images restricted to a single frame in the z axis that lies in the upper 1/3 of the majority of OHCs and shows colocalization of prestin and αII spectrin in the lateral wall of OHCs. The upper panel shows the corresponding z plane reconstruction of a plane vertical to OHCs (light blue line). It shows αII spectrin colocalizing with prestin in the lateral wall and also densely expressed in the cuticular plate at the apical end (where prestin is absent/ minimal). Scale bar = 20 mm. G-I: Confocal image of a row of OHC and underlying Deiter cells viewed laterally shows spectrin immunolabel in the lateral wall of OHCs along with phalloidin labeled b actin. The apical surface of OHCs show dense actin and spectrin label in the cuticular plate and actin rich stereocilia overlying the cuticular plate (Scale bar = 5 mm).

We then used super resolution microscopy to localize spectrin in the lateral wall of OHCs. Figures 2, 3 and 4 show STORM (2,3) and STED (4) images of OHCs immunolabeled with antibodies to αΙΙ spectrin and prestin. We observe that the two proteins are sharply colocalized. Using the nearest neighbor algorithm to ascertain distances between the two labels, we find the distribution to be right skewed with the mode in the distances between αΙΙ spectrin and prestin to be 20-25nm (Figure 2). These findings are in approximation to distances between the plasma membrane and spectrins calculated by other investigators using electron microscopy (*13*). When viewed in the axial and coronal planes we also observe an unexpected periodicity in the signal intensity of αΙΙ spectrin. In both planes these clusters had variable periodicity. We did not have sufficient data (in the order of 1000s of frames) to make a valid statistical model. However, when the z plane was restricted to 500nm the shortest distance between the centers of adjacent clusters was in the order of ∼ 3-400 nm. The clusters themselves were as small as 150nm and as large as 800 nm in the XY plane. Note that lacking a reference frame, we are unable to correct tilts of the cell in the vertical plane. We observe a similar periodicity when images were acquired with a STED microscope in the axial plane, where the distances from peak to peak varied from 350nm to 700nm. Since the images are of axial sections of OHCs where spectrin filaments are arranged orthogonally, these densities likely represent spectrin filaments that are either in register or as local collections. The diameter and periodicity are too large to be that of individual filaments (*30, 31*). These data will be explored in detail in a further manuscript on the role beta spectrins in OHC function (Bai et al., in preparation).

**Figure 2.**
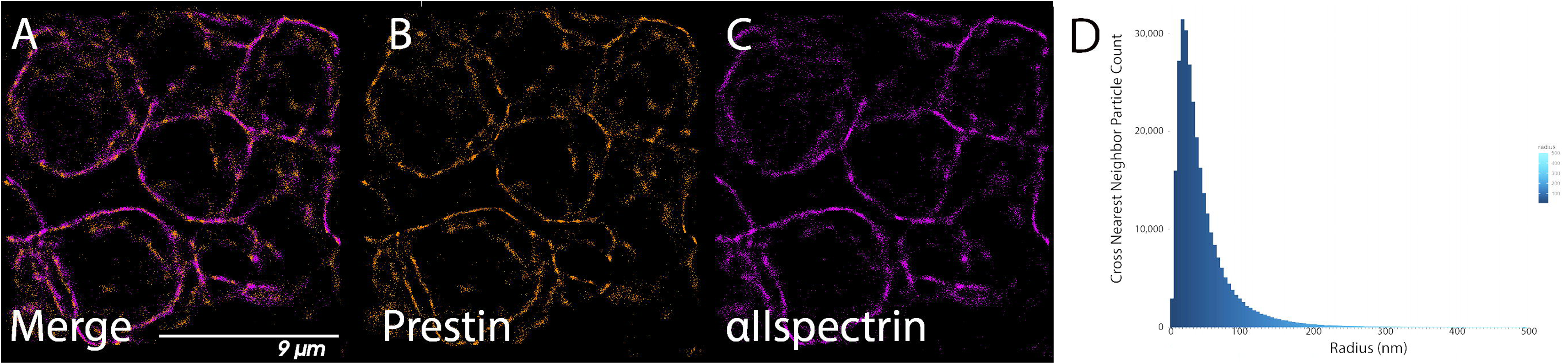
Shown are pointillistic based STORM images of OHCs immunolabeled with prestin (B) and αII spectrin (C) with merged images in A. Hair cells (4+) are seen in axial views. The z depth is restricted to 1mm with undulating OHC plasma membranes evident in the images. The two labels are sharply colocalized (merged image A) with the nearest neighbor algorithm showing particles with a right skew in distribution (D) with the mode in distribution to be 20-25nm. Other co-localization metrics include Li’s intensity correlation of 0.78 and Pearsons correlation of 0.5.

**Figure 3.**
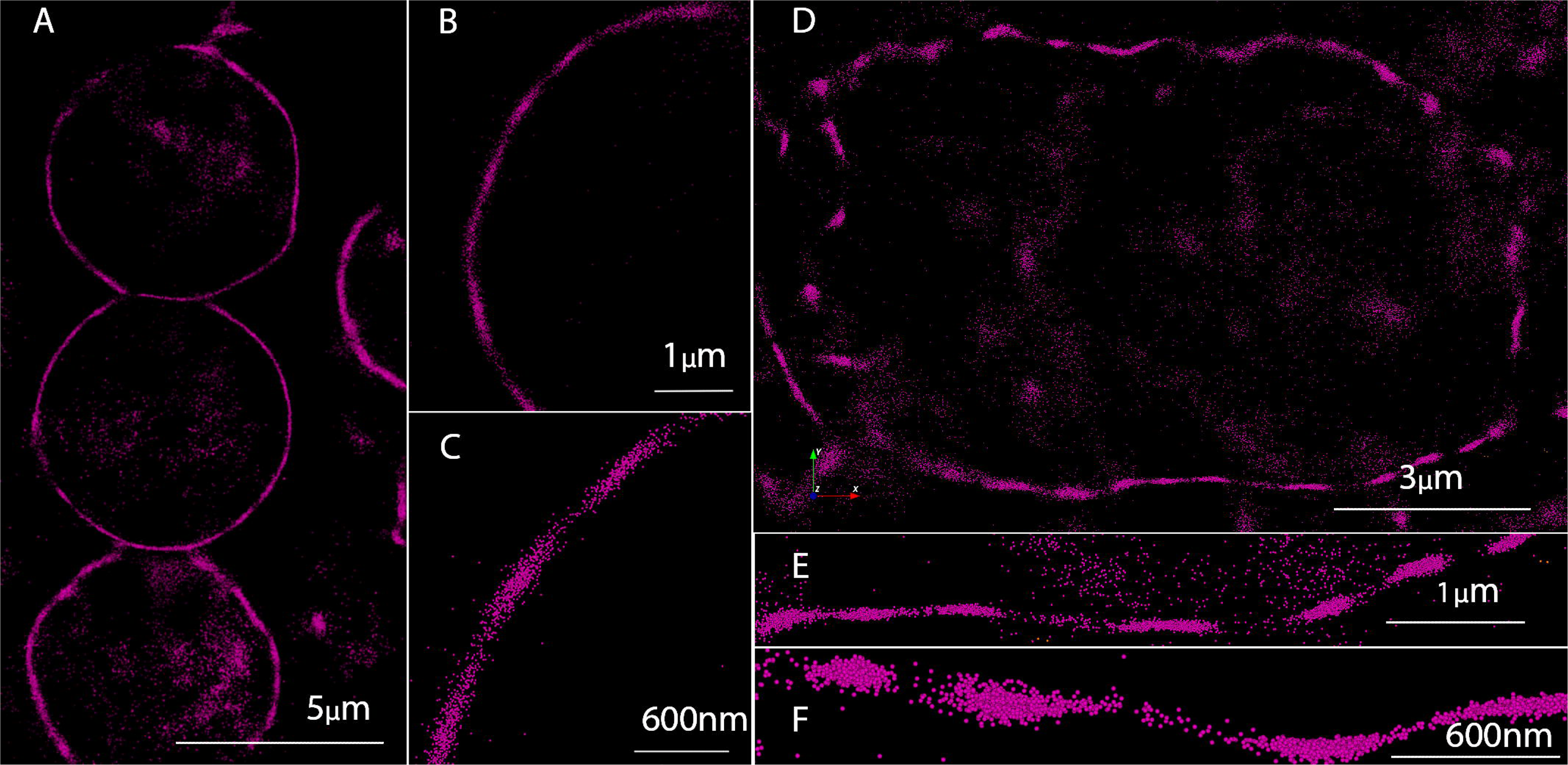
STORM images show clustering in the expression of αII spectrin detected by immunolabeling along the lateral wall of the cell in the axial and more coronal planes. Depicted are representative images in the axial (A-C) and more coronal (D-F) planes with each series increasing in magnification and from different areas of the lateral wall of cells. The size of the clusters and the distance between them varied. The clusters were 80 to 800 nm in size. The intercluster distances were 150 to 750 nm in size. It was our impression that the clusters evident in the coronal plane were smaller but do not have sufficient data (>100 images) to make a confident statement.

**Figure 4.**
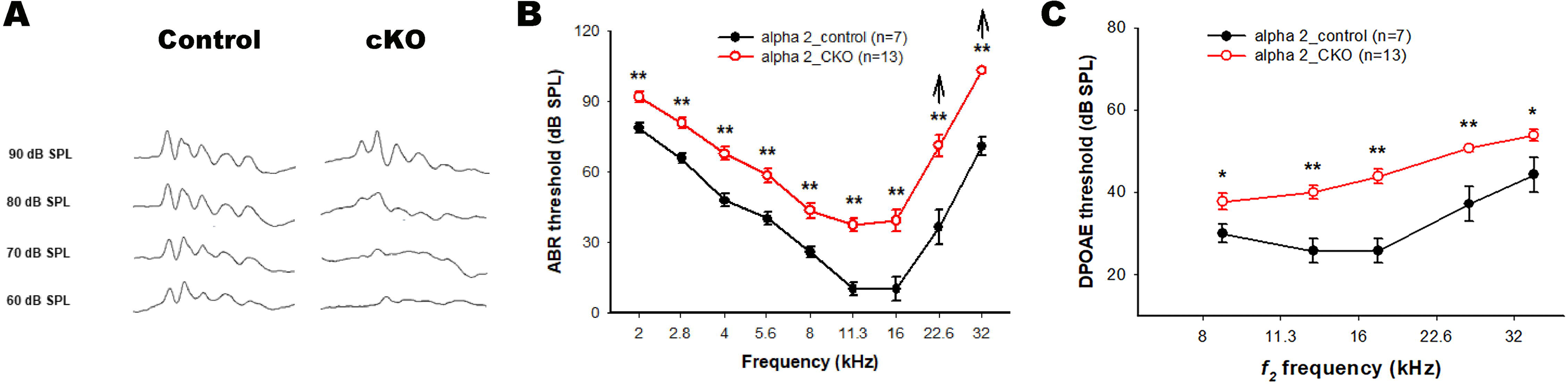
STED images confirm STORM imaging showing αII spectrin at the lateral wall of OHCs where it co-localizes with actin. Shown are oblique axial views of two rows of OHC with the lateral wall in the lower half of each OHC and the cuticular plate in the upper half. These sections were taken below the cuticular plate and contain a significant amount of actin and αII spectrin extending below the cuticular plate in the cytoplasm. Phalloidin labeled actin (B) lies in close approximation with immunolocalized αII spectrin (C) with the merged image in A. The inset in C shows a profile of αII spectrin intensity in the lateral wall of an OHC (boxed area in C). Here too, we note a periodicity in the intensity of staining that is similar to that of αII spectrin seen on STORM imaging. To allow clearer visualization we present panels B and C in black and white. Scale bar = 3 mm.

### Conditional knockouts of αΙΙ spectrin in the postnatal mouse result in loss of αΙΙ spectrin from the lateral wall with preserved OHCs including their stereocilia

To test the effects of deleting αΙΙ spectrin in mouse OHCs, we used prestin Cre mice crossed with Spatn1 fl/fl mice and induced cre by injecting tamoxifen between post-natal days 9-15. In this way we were able to specifically knockout αΙΙ spectrin in OHCs alone without affecting IHCs. In addition, we were able to knockout αΙΙ spectrin in these cells at a later time point that was closer to their functional maturity. Thus, we avoided the large-scale loss of these cells that were observed in the two previous knockout models using GfiCre and Pou4f3Cre that resulted in a knockout of αΙΙ spectrin and βΙΙ spectrin as early as E16.5 and E14.5 respectively. In the case of the βΙΙ spectrin knockout, there was no detectable αΙΙ spectrin in the cuticular plate region of these mice.

We make several observations. 1. Knockdown of αΙΙ spectrin was observed in most OHCs. 2. αΙΙ spectrin was consistently absent in the lateral wall of OHCs. 3. Low levels of αΙΙ spectrin immunofluorescence in cuticular plates was observed even as late as P90 suggesting that the protein undergoes slow turnover. 4. Concordant with this finding, stereocilia were observed in most OHC. 5. OHC loss on the one hand and lack of αΙΙ spectrin knockdown on the other was sporadic. Figure 5 shows a representative confocal image of the Organ of Corti from a Prestin Cre+ spectrin fl/fl mouse that was induced with tamoxifen between P9 and P15. The mice tissue were harvested at P90. These images were taken from the apical turn of the cochlea and shows preserved IHCs and most OHCs. In addition, there is a loss of αΙΙ spectrin immunolabeling from the lateral wall of these cells. We also note a diminished staining of αΙΙ spectrin in the cuticular plate of these OHCs and particularly when compared to IHC where the staining is normally equivalent.

**Figure 5.**
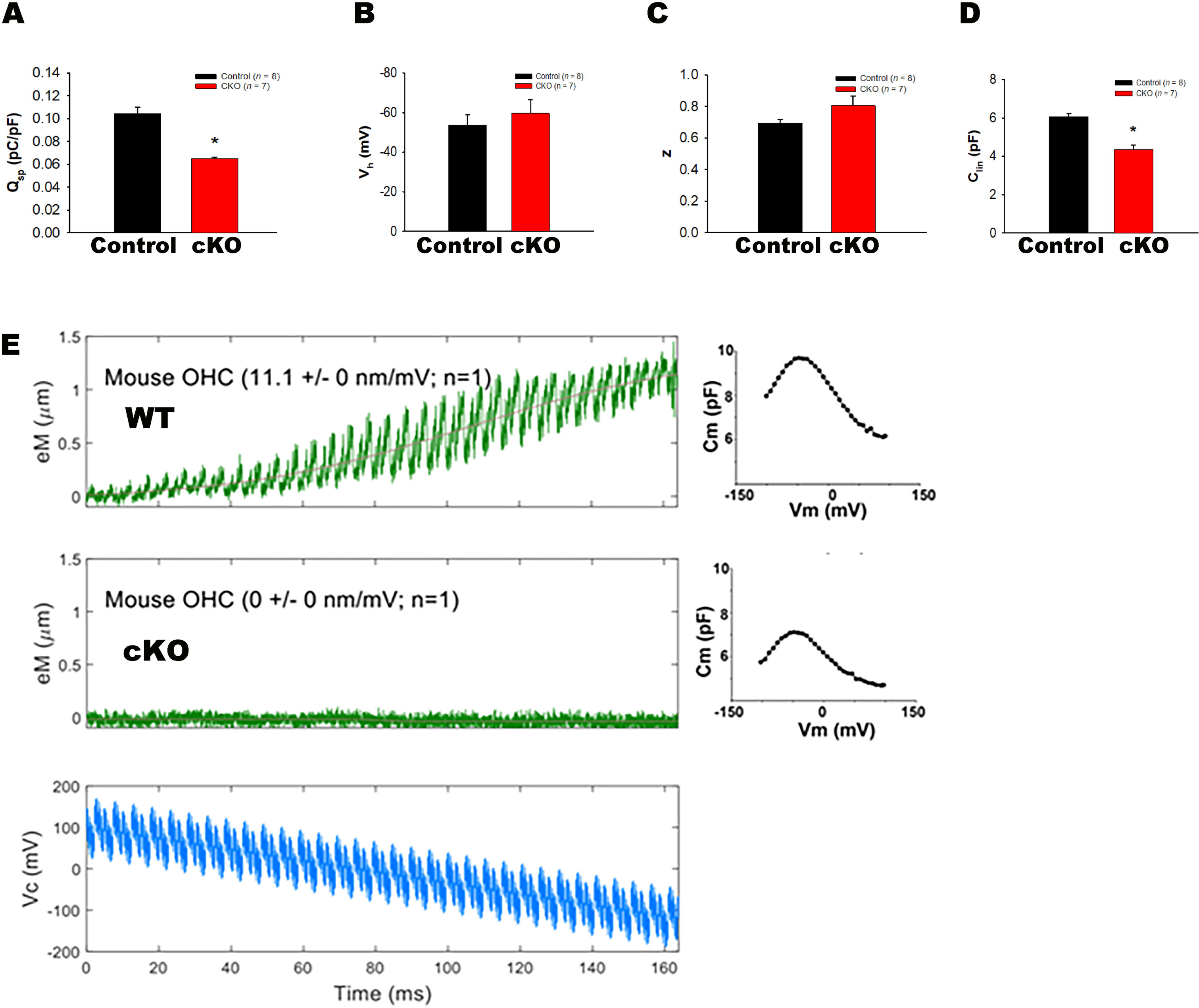
Shown are confocal images of OHCs from αII spectrin knockout mice that were immunofluorescence labeled with antibodies to αII spectrin and actin labeled with phalloidin Alexa 647 (A-I). A-C are 3D projections (maximum intensity) of a z stack confocal image OHCs and IHCs in which actin (B) and αII spectrin (C) were detected by (immuno) fluorescence (A, merged). Note the significant loss of αII spectrin staining in the OHCs when compared to IHCs (compare these to WT animals in Figure 1). D-I. Shown are confocal images taken through a single z plane of the same image showing stereociliary bundles labeled with phalloidin (E, H) from the OHCs of row 2 (D-F) and row 3 (G-I) with concordant αII spectrin label (F, I). Merged images are in D and G). J-L are lateral views images of OHCs labeled with antibodies to prestin (K) and αII spectrin (L) showing knockdown and especially in the lateral wall. Merged images in J.

### Ultrastructural changes in OHCs lateral wall have parallels seen in neuronal cells of αΙΙ spectrin knockouts

We examined the ultrastructure of OHCs from knockout mice using transmission electron microscopy. Additionally, we also performed tomography from tilt series of several OHCs that also allowed us to obtain volumetric information of the lateral wall. We made several observations (**Figure 6**) in the αΙΙ spectrin cKO mice: 1 The volume between the plasma membrane and the outer membrane of the subsurface cisternae was increased. 2. The widths and volumes of the subsurface cisternae were increased. 3. In places we observed two layers of the subsurface cisternae. In contrast, WT mice consistently demonstrate a single subsurface cisternal layer. 4. In reconstructions of the sub-plasma membrane space, we noted a significant decrease in filament density. Of note, we did not find alterations in the size/ density of the cuticular plates or the overlying stereocilia (Figure 6).

**Figure 6.**
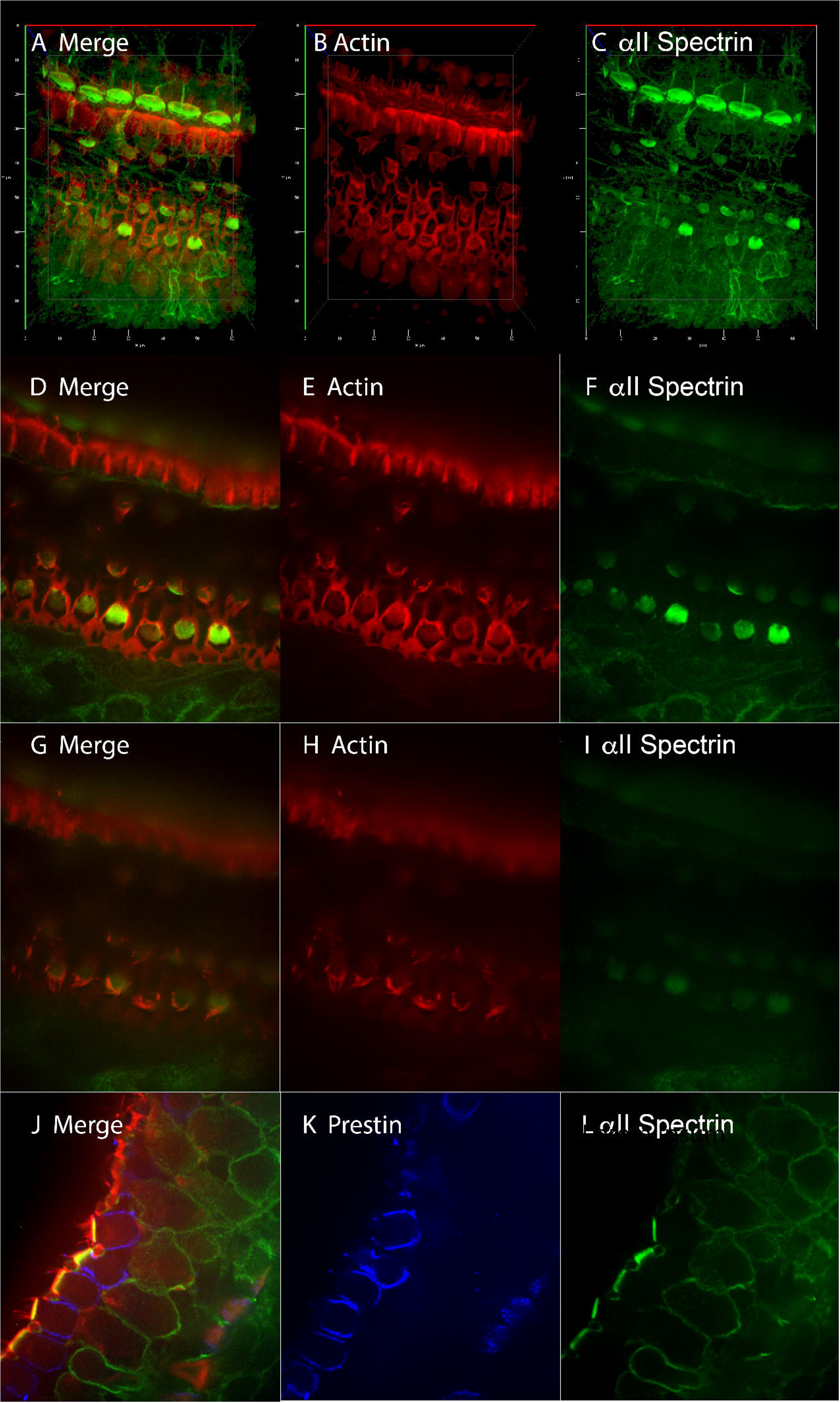
Ultrastructural changes in the lateral wall of αII spectrin cKO mice observed on transmission electron microscopy. **A.** Shown is a single OHC with areas where the SSC were duplicated (*). Two of these areas (arrow) are shown at higher magnification. The cell also has an intact cuticular plate and overlying stereocilia. **B, C.** A tilt series and reconstructions from WT (B) and cKO mice is shown. The upper left panel is the X-Y depiction of the electron micrographs. The upper right panel shows the YZ projection at the yellow + sign. The lower panels are the XZ projections and show a network of electron dense filaments traversing the space. These networks are significantly diminished in the αII spectrin cKO mice (**C**) lower panel compared to WT (**B**). **D,E**. Shown are X-Z reconstructions with segmentation of the plasma membrane (green) and sub-surface cisternae (SSC (cyan)) of WT (**D**) and αII spectrin cKO mice (**E**). **F, G**. The volumes of the SSC and the space between the plasma membrane and SSC are both significantly enlarged in a spectrin cKO mice (SSC volume per 1000 planes: WT: 4426122±518954.3 nm^3^ ; αII spectrin cKO: 7696776.67±773127.2 nm^3^; p= 0.014 on a students t test. PM-SSC volume per 300 planes: WT: 2099277.5±59745.5 nm^3^; αII spectrin cKO 2439280± 84361.8 nm^3^. P= 0.019 on a students t test.

### The auditory phenotype in αΙΙ spectrin indicate OHC dysfunction

We determined the auditory phenotype of cKO mice with ABRs and DPOAE. Tamoxifen injections produced variable hearing loss in αΙΙ spectrin cKO mice with a bimodal distribution in thresholds. 80% of αΙΙ spectrin cKO mice showed an increase in hearing thresholds after tamoxifen induction. Approximately 20% of mice showed minimal hearing loss. OHCs from these mice showed no loss of spectrin expression on confocal immunofluorescence. In contrast, we observed a marked reduction in OHC spectrin in those mice that showed a significant hearing loss. We attributed this discrepancy to the inefficient induction of Cre using the prestin-Cre model, an effect that has been previously observed (*32*). When we limited our analysis to mice with a demonstrable hearing loss, we found a 20 dB increase in thresholds at 16kHz that also extended across the frequency spectrum from 2kHz to 32kHz (**Figure 7B and C**). There was a slight further increase in the drop-off in hearing sensitivity at higher frequencies and a slight decrease in the drop-off in hearing sensitivity at lower frequencies. DPOAE thresholds also showed an ∼ 20dB increase in thresholds in αΙΙ spectrin cKO mice (**Figure 7C**). Here too, we limited our analysis to those mice that showed an increase in ABR thresholds. As with ABR threshold increases, the increase in DPOAE thresholds extended across the frequency spectrum.

**Figure 7.**
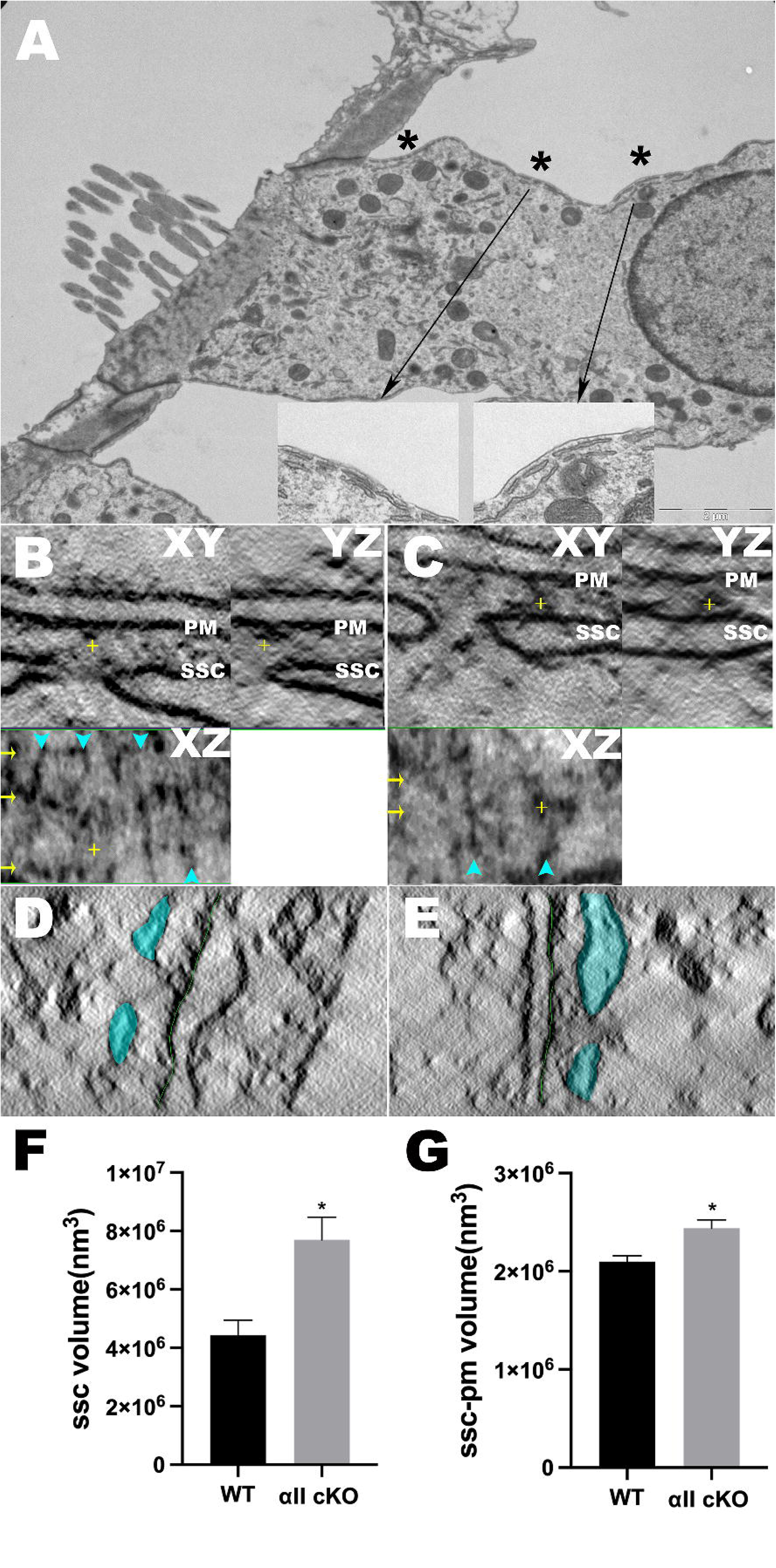
αII spectrin cKO mice have an auditory phenotype that confirms OHC dysfunction. A. Shown are representative ABR traces at 16kHz from WT and αII spectrin cKO mice at different intensities of sound. B. Mean ABR thresholds (+/-SEM) of αII spectrin cKO mice (red) are shown across the frequency spectrum and compared to WT mice (black). There is an increase in thresholds of 20dB in cKO mice at 16kHz. There is a further increase in the thresholds to 35dB at 22 and 32 kHz where OHC function is thought to be more critical. C. DPOAE responses of cKO mice (red) demonstrate an increase in thresholds of ∼ 20dB compared to WT mice (black).

### There is an uncoupling of electromotility from prestins gating charge movement in OHC of αΙΙ spectrin cKO mice

We measured prestins gating charge movement determined as a voltage dependent increase in non-linear capacitance. The most notable change we found in αΙΙ spectrin cKO mice was a decrease in prestin density (Figure 8A) evidenced by a 40% decrease in Q_sp_ (total gating charge per unit of linear capacitance). Importantly, we observed a significant 25% decrease in the linear capacitance of these cells indicating that the cells were also smaller (**Figure 8F**). We think the alternative that their lipid composition and membrane thickness was altered to be unlikely, since our measurements of OHC lengths were consistent (with smaller cells). There were two other findings including a slight 5mV hyperpolarizing shift in the voltage at peak capacitance (V_h_) (8B) and a 15% *increase* in our estimate of the unitary charge movement z (8C).

**Figure 8.**
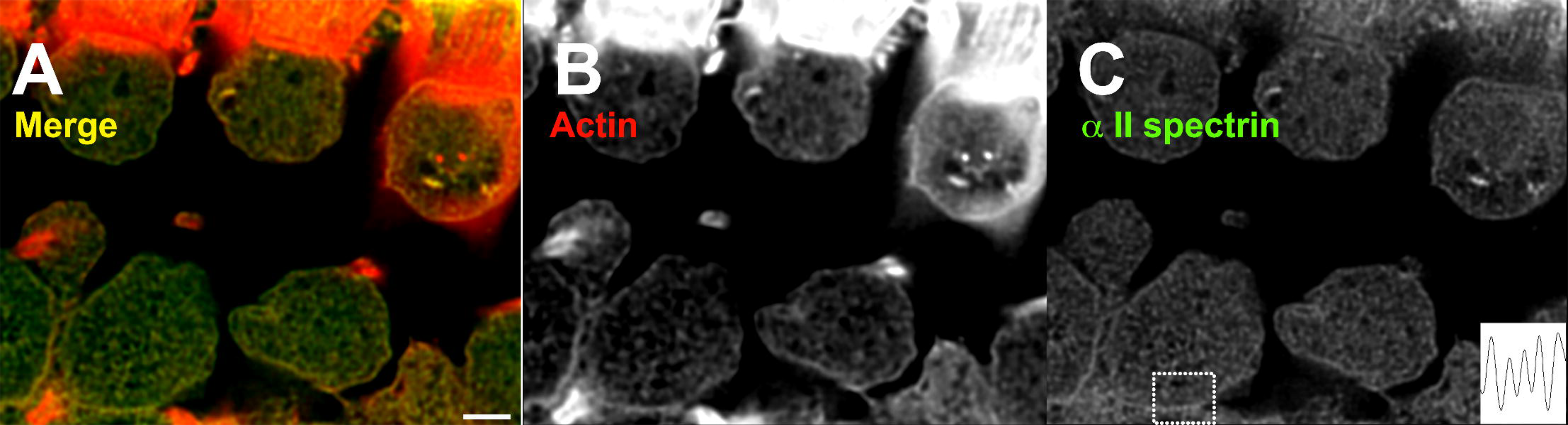
Electromotility and prestins gating charge movement can be disassociated in OHC of αII spectrin cKO mice. **A-D.** Shown are measures of NLC from OHC of αII spectrin cKO mice (red) compared to WT control mice. OHC of αII spectrin cKO mice show a 38% reduction in Q_sp_ (**A**), a measure of prestin density in the membrane compared to WT mice (WT 0.104 +/- 0.005 pC/pF (n=8); cKO 0.065 +/- 0.008 pC/pF SEM (n=7); P<0.05 in a students t test). These cKO mice OHCs also show a slight ∼5mV hyperpolarizing shift in V_h_ (**B**) (WT-53.724 +/- 5.091 mV (n=8); cKO -59.515 +/- 6.947 mV (n=7)) and a 16% increase in our estimate of unitary charge z (**C**) (WT 0.692 +/- 0.024 e (n=8); cKO 0.806 +/- 0.06e (n=7); p> 0.05). OHC of αII spectrin cKO mice also show a 20% reduction in the linear capacitance (WT 6.06 +/- 0.15pF (n=8); cKO 4.34 +/- 0.24pF (n=7); P<0.05 in a students t test (D)). E. Shown are electromotile responses to a voltage ramp of a single representative WT (upper panel) OHC compared to cKO OHC (middle panel). The lowest panel shows the corresponding transmembrane voltage and time series. We also include NLC traces from the same OHCs from WT (upper right) and cKO mice (lower right). Averages for electromotility were significantly different (WT 9.7 +/- 0.9 nm/mv SEM (n=8); cKO 1.8+/- 0.59 nm/mV SEM (n=6). P<0.05 in a students t test).

We measured the electromotile response of OHCs in response to a change in voltage as previously described (*33*). Here, we found OHCs of αΙΙ spectrin cKO to demonstrate an ∼ 80% reduction in electromotility. The ∼ 80% reduction in electromotility was larger than the reduction in total prestin associated gating charge movement (60%). As noted previously, Q_sp_ the measure of gating charge movement per unit of membrane was reduced by 38%. Notably, a few cells demonstrated no eM while having clearly measurable NLC (**Figure 8D**).

## Discussion

In this paper we describe the effects of targeted deletion of αΙΙ spectrin in OHCs in the post-natal period that causes a loss in electromotility and resultant diminished hearing. These data stand in contrast to previous experiments that targeted αΙΙ spectrin from both inner and outer hair cells in the embryonic period that caused variable loss of hair cells, disordered stereocilia and deafness. The key finding in our data is that in the postnatal OHC loss of spectrin affects electromotility.

Previous experiments have confirmed the likely presence of αΙΙ spectrin in the lateral wall of OHCs. Our use of knockout validated antibodies that confirmed the presence of the protein in the lateral wall solidifies these findings further. We present here a plethora of data substantiating dysfunction in the lateral wall of OHCs. A key observation on both confocal microscopy and transmission electron microscopy was that the cuticular plate and overlying stereocilia were not affected by the deletion of αΙΙ spectrin in the postnatal period. This may speak to the need of the protein in these structures in developing hair cells since a prominent defect was the loss of stereocilia in embryonic knockouts.

On superresolution STORM microscopy we were able to assess the distances between immunolocalized prestin and αΙΙ spectrin molecules. We see a skewed distribution in the distances with the mode of the distances lying between 20 and 25nm. These values are consistent with previous estimates of the distance between the plasma membrane, where prestin resides, and the underlying spectrin cytoskeleton (*13*). We also observed a periodicity in spectrin on super-resolution microscopy using STORM and STED microscopy that has a resolution limit of 25-50nm. In our experiments, the observed periodicity varied significantly with some in the range previously described in neurons (180nm) and consistent with an extended form of spectrin (*30, 31*). In our limited dataset, a significant number of the intercluster distances was in the vicinity of 500nm. These data will be explored in detail in our paper on the effects of beta spectrin knockouts on OHC function (Bai et al., manuscript in preparation).

Our EM data show consistently enlarged subsurface cisternae and the space between the subsurface cisternae and the plasma membrane in αΙΙ spectrin cKO OHCs. The physiological significance of enlarging these spaces is unclear, although the space between the plasma membrane and SSC has been implicated in maintaining chloride concentrations and even as an electrically isolated compartment (*34, 35*). It should be noted that two groups have reported the presence of a dense cytoskeletal network adjacent to the plasma membrane, and also to the subsurface cisternae (*14, 15*). Hence the changes in these volumes by disruption of spectrin networks could be anticipated.

We observed a bimodal hearing phenotype in our tamoxifen induced mice with ∼ 80% of mice showing a hearing phenotype and the remainder showing no hearing loss. We attributed the differences in hearing loss to an incomplete induction by tamoxifen in the prestin-Cre mice in the minority that did not show hearing loss. Our immunofluorescence data showing spectrin expression in OHCs of mice with no phenotype and loss of OHC spectrin in mice with a phenotype substantiates this hypothesis. Moreover, the incomplete knockdown using the tdTomato reporter has also been observed in other experiments where prestin Cre mice was used (*32*).

On measuring NLC, we observe a 38% reduction in Q_sp_; Q_sp_ is a measure of prestin density in the membrane in OHCs of αΙΙ spectrin cKO. These data suggest a role for spectrin in movement of prestin from the ER to the plasma membrane or its removal or even in reducing prestin transcription. Irrespective, it is unlikely that the reduced prestin expression has a significant effect on hearing thresholds or DPOAEs since previous data have shown normal hearing when prestin expression was by 66% (*36*).

We note that αΙΙ spectrin cKO mice show a minimal 5mV hyperpolarizing shift in V_h_. These data are an indirect measure of normal membrane tension in αΙΙ spectrin OHCs. Similarly, the tendency for increased z (that was however, not statistically significant) could represent a loss of constraints from tethering of prestin to the underlying cytoskeleton.

Our experiments confirm a disproportionate loss of electromotility in OHCs of αΙΙ spectrin cKO that cannot be explained by changes in prestin expression in OHCs. In some cells, there was measurable NLC and yet absent electromotility. Previously, it was hypothesized that electromotility was a membrane-based phenomenon, with differences in axial and circumferential tension directing the force generated by prestins expansion in a longitudinal / axial direction. In this formulation, the role of the cytoskeleton was largely passive serving to generate differences in axial and circumferential tension. Our data showing discordant loss of electromotility in αΙΙ spectrin mice suggests a more active role for the cytoskeleton in force transmission during electromotility as had been shown with in vitro with diamide treatment. In this context, it is important to note that minimal changes in V_h_ suggests that tension/ load changes are not a significant factor in the changes we observe. We conclude that αΙΙ spectrin is critical for force transmission in electromotility and its loss is the chief cause of hearing loss in αΙΙ spectrin cKO mice. These data further cement the molecular identity of spectrins in the lateral wall (*37*) and establish their physiological role in the lateral wall of OHCs.

## Methods

### Animals

We used αII spectrin floxed mice (*Sptan1^(^*^FF)^) that were a gift from the Rasband lab (*38*). We then crossed these mice with prestin-CreERT2 mice (a gift from the Zuo lab (*39*)) that allowed an inducible postnatal OHC specific knockdown of αII spectrin (outlined in Supplementary Figure 1). Briefly, αII spectrin floxed mice and control mice were injected with tamoxifen at 1-2 week (between p9-p15). The concentration of tamoxifen used was calculated based on mouse body weight (4 mg/40g body weight). A stock solution [10mg/ml] of tamoxifen was prepared in corn oil and kept at 4°C for up to 2 weeks. A 30G syringe was used to inject tamoxifen into the peritoneum (I.P.). Doses were administered on three consecutive days and to minimize distress to the animal, the side of injection is alternated day-by day. For genotyping animals, ear punches were collected from mice at their weaning age. Genomic DNA was prepared by an overnight 56°C incubation in lysis buffer 50mM Tris pH 8.0, 50mM KCL, 2.5mM EDTA, 0.4% Igepal, 0.4% Tween 20 supplemented with Proteinase K (400 μg/ml). gDNA was assayed by PCR using the following primer sets:

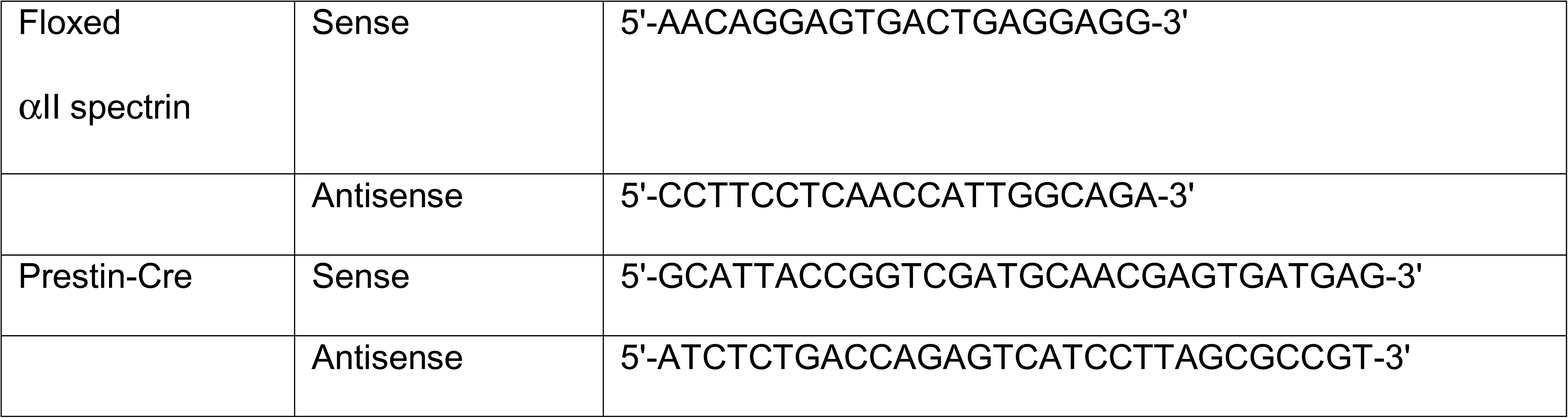

All animal experiments were performed according to a protocol approved by the Yale University Institutional Animal Care and Use Committee (IACUC) and animals were cared for in accordance with the recommendations in the “Guide for the Care and Use of Laboratory Animals” (National Institutes of Health). Both WT and cKO mice were raised in the same room within the Yale Animal Resources Center (YARC). Mice were housed in standard cages and maintained on a 12 h light–dark cycle. They had ad libitum access to drinking water and normal diet throughout the experiment. Mice of both genders were used in the study.

### Confocal Immunofluorescence

Immunofluorescence staining of freshly isolated apical turns and fixed and decalcified three turns of the organ of Corti were performed as previously described (*40, 41*). Apical turns of the organ of Corti were dissected and fixed in 4% PFA for 30 minutes, washed in PBS 0.05% Tween 20, 0.05% Triton-X 100 (incubation buffer) 3 times of 3 minutes each, and incubated with primary antibody in incubation buffer overnight at 4°C. The primary antibodies used included goat anti-prestin (1:500; Santa Cruz, cat # sc-22692), mouse anti-αII spectrin (1:200; Biolegend, cat # AG 803206). After 3 washes in incubation buffer the tissue was incubated with secondary antibody in incubation buffer for 1 hour at room temperature. The secondary antibodies used included donkey anti-mouse Alexa 488 (1:200; Invitrogen, cat # A-21202), donkey anti-goat Alexa 647 (1:200; Invitrogen, cat # A32723). Actin was detected using Phalloidin Alexa Fluor 647 (1:200; Abnova, cat # U0298) added to the secondary antibody step. Tissue was then washed thrice in incubation buffer and then mounted in Vectashield Antifade Mounting Medium (Vector Laboratories, Inc., Burlingame, CA, USA) and viewed using a Zeiss spinning disc laser scanning microscope (Zeiss Observer Z1).

### Direct Stochastic Optical Reconstruction Microscopy (dSTORM)

Super-resolution dSTORM imaging was performed as previously described (*42*). In brief, the apical turn of the organ of Corti was freshly dissected and hair cells exposed by removal of the tectorial membrane. Tissue was pre-extracted with 0.2% saponin for 1 minute followed by fixation with 3% PFA and 0.1% glutaraldehyde for 15 minutes. The tissue was reduced with 0.1% NaBH_4_ for 7 minutes and then labeled with primary antibodies (goat anti-prestin, 1:100; mouse anti-αII spectrin, 1:200; Phalloidin Alexa Fluor 647, 1:200) for 1 hour and secondary antibodies (donkey anti-mouse CF 568, donkey anti-goat Alexa 647; 1:500) for 45 minutes after blocking, with 3 washes of 3 minutes each between each step. The sample was post-fixed after antibody labeling with 4% PFA for 10 minutes. Freshly made imaging buffer containing glucose oxidase (135 U/mL), catalase (1,000 U/mL), β-mercaptoethanol (1% v/v), and MEA (20 mM) was added just before imaging. Super-resolution dSTORM images were obtained with a Bruker Vutara 352 Super-Resolution Microscope (Bruker Nano Surfaces, Middleton, WI, USA) with a 60X 1.2 NA objective and a 1 W 561 nm and 640 nm laser. Imaging beads confirmed that resolution was 20 nm in the *xy* plane and 50 nm in the *z*-direction. Calibration before experimentation was done by calculating the point spread function (PSF) in three dimensions using beads. Images were rendered and analyzed with Vutara’s SRX localization and visualization software (v6.2). Images were obtained in both planes simultaneously. The background was removed after the frames were obtained and particles identified on their brightness. Three-dimensional localization of the particles was based on the 3D model function that was obtained from recorded bead datasets. The recorded fields were aligned automatically by computing the offline transformation between the pair of planes. Typically, we collected 5000 frames with each fluorophore using 20 µsec times. Data were analyzed using algorithms embedded in the Vutura software (v6.2). These include the Crossed Nearest Neighbor algorithm and cluster identification. All chemicals were purchased from Sigma-Aldrich.

### Gated Stimulated Emission Depletion Microscopy (gSTED)

STED images were acquired using a Leica SP8 3X STED using a high-contrast Plan Apochromat 100× 1.40 NA oil CS2 objective (LAS X; Leica Microsystems). Samples were prepared as for STORM imaging with the following modifications. Primary antibody was used at 1:50 and secondary antibody (Alexa 555) at 1:100 dilutions. Phaloidin Alexa 647 was used at 1:50 dilution. Samples on Titertek 1.5 glass dishes were embedded in Prolong Diamond (Invitrogen) overnight before imaging. Depletion and acquisition of Alexa 647 images was done prior to Alexa 555. Deconvolution of the acquired datasets was done posthoc using Huygens software.

### Auditory Brainstem Responses (ABR) and Distortion Product Otoacoustic Emissions (DPOAE)

Hearing sensitivity of WT and a a spectrin cKO mice was measured using two routinely used non-invasive hearing tests: ABRs, which represent synchronized electrical activity in the auditory nerve and ascending central auditory pathways in response to acoustic stimuli, and DPOAEs, which are sounds generated by cochlear OHCs in response to a two-tone stimulus. Measurements were carried out within a sound attenuating booth (Industrial Acoustics Corp., Bronx, NY, USA). Mice were anaesthetized with chloral hydrate (480 mg/kg), injected intraperitoneally (IP), and placed onto a heating blanket (Harvard Animal Blanket Control Unit, Harvard Apparatus Ltd., Kent, England), to maintain body temperature at 37°C. Prior to measurement, an otoscopic examination of the tympanic membranes were performed on the anesthetized mouse using an ENT surgical microscope (ZEISS OPMI 1-FC, Carl Zeiss, Oberkochen, Germany) for evidence of otitis media (middle ear infection). Mice displaying signs of otitis media in either or both ears were excluded from the study. The acoustic stimuli for ABR and DPOAE were produced and the responses recorded using a TDT System 3 (Tucker-Davis Technologies, Inc., Alachua, FL, USA) controlled by BioSigRP (TDT), a digital signal processing software.

ABR measurements were performed as previously described (*43*) by placing subdermal needle electrodes (LifeSync Neuro, Coral Springs, FL, USA) at the vertex (active, noninverting), the right infra-auricular mastoid region (reference, inverting), and the left neck region (ground). ABRs were elicited with pure tone pips presented free field via a speaker (EC1 Electrostatic Speaker, TDT) positioned 10 cm from the vertex. Symmetrically shaped tone bursts were 3 ms long (1 ms raised cosine on/off ramps and 1 ms plateau) and were delivered at a rate of 21 per second. Stimuli were presented at frequencies between 2 and 32 kHz in half octave steps and in 5 decibel (dB) decrements of sound intensity from 90 dB SPL (sound pressure level) (or 110 dB SPL if thresholds exceeded 90 dB SPL). Differentially recorded scalp potentials were bandpass filtered between 0.05 and 3 kHz over a 15-ms epoch. A total of 400 responses were averaged for each waveform for each stimulus condition. The ABR threshold was defined as the lowest sound intensity capable of evoking a reproducible, visually detectable response. Suprathreshold amplitudes (µV) and latencies (ms) of the initial four ABR waves (waves I, II, III and IV) were then determined at 16 kHz. The most sensitive frequency range of hearing in mice is 11.3 to 22.6 kHz, and 16 kHz is half octave in-between so was therefore chosen for analysis. The analysis was carried out offline in BioSigRP on traces with visible peaks by setting cursors at the maxima and minima (trough) of the peaks. Latency was determined as the time from the onset of the stimulus to the peak while amplitude was measured by taking the mean of the ΔV of the upward and downward slopes of the peak.

DPOAEs were measured by inserting a microphone probe (ER-10B+ Microphone System, Etymotic Research, Inc., Elk Grove Village, IL, USA) into the external ear canal of the anesthetized mouse, with two speakers (MF1 Multi-Field Magnetic Speakers, TDT) connected to the probe via tubing. Two simultaneous continuous pure tones (*f*_1_ and *f*_2_) that have a frequency ratio of 1.2 (*f*_2_/*f*_1_) and equal sound level (*L*_1_ = *L*_2_) were delivered at center frequencies of 8, 12, 16, 24 and 32 kHz with sound levels from 80 to 20 dB SPL in 10 dB decrements. Stimulus duration was 83.88 ms at a repetition rate of 11.92 per second. The acquired DPOAE responses were averaged 100 times. The DPOAE of interest was at 2*f*_1_-*f*_2_, the largest and most prominent DPOAE. The DPOAE threshold was defined as the lowest sound intensity capable of evoking a visually detectable 2*f*_1_-*f*_2_ signal above the noise floor. DPOAE amplitudes were analyzed offline in BioSigRP by setting cursors at the peak of the 2*f*_1_-*f*_2_ signals.

### Non-Linear Capacitance (NLC)

NLC was recorded in OHCs from WT and KO mice as previously described (*44*). Mice were euthanized with an overdose of chloral hydrate (480 mg/Kg; IP), the cochleae extracted, and the apical turn of the organ of Corti dissected out. The cochlear explant was recorded with ionic current blocking solutions, thereby removing interference on measures of membrane capacitance. The extracellular solution contained (in mM): 100 NaCl, 20 tetraethylammonium (TEA)-Cl, 20 CsCl, 2 CoCl_2_, 1 MgCl_2_, 1 CaCl_2_, 10 HEPES, pH 7.2. The intracellular solution contained (in mM): 140 CsCl, 2 MgCl_2_, 10 HEPES, and 10 EGTA, pH 7.2. Patch pipettes had resistances of 3-5 MΩ. Gigaohm seals were made and stray capacitance was balanced out with amplifier circuitry prior to establishing whole-cell conditions. A Nikon Eclipse E600-FN microscope (Nikon Instruments, Inc., Melville, NY, USA) with 40X water immersion lens was used to observe cells during voltage clamp. Whole cell voltage clamp recordings were performed from all rows of OHCs of the whole mount organ of Corti. OHCs were recorded at room temperature using jClamp software (www.scisoftco.com) and an Axopatch 200B amplifier (Molecular Devices, LLC, San Jose, CA, USA). Data were low pass filtered at 10 kHz and digitized at 100 kHz with a Digidata 1320A (Molecular Devices, LLC). In order to extract Boltzmann parameters, capacitance-voltage data were fit to the first derivative of a two-state Boltzmann function.

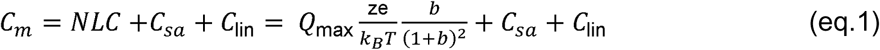

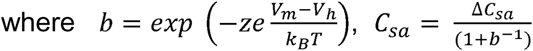

Q_max_ is the maximum nonlinear charge moved, V_h_ is voltage at peak capacitance or equivalently, at half-maximum charge transfer, V_m_ is R_s_-corrected membrane potential, z is valence, C_lin_ is linear membrane capacitance, e is electron charge, *k_B_* is Boltzmann’s constant, and T is absolute temperature.

### Transmission Electron Microscopy (TEM)

Morphology of OHCs from WT and αII spectrin cKO mice were examined using TEM. Mice were euthanized with an overdose of chloral hydrate (480 mg/Kg; IP), the cochleae extracted, then fixed with 2% paraformaldehyde and 2.5% glutaraldehyde in 0.1 M sodium cacodylate buffer (pH 7.4) for 30 minutes at room temperature and then overnight at 4°C (cochleae were perfused through the round and oval windows, and then immersed in the fixative solution). After fixation, cochleae were washed three times with sodium cacodylate buffer, and then the apical turn of the organ of Corti was dissected out. Subsequently, samples were post-fixed in 1% osmium tetroxide in 0.1 M sodium cacodylate for 30 minutes, followed by dehydration in a graded ethanol series (50%, 60%, 70%, 80%, 90%, 100%). After replacement with a combination of propylene oxide and Eponate 12^TM^ Embedding kits (Ted Pella, Inc., Redding, CA, USA), the samples were embedded in the module and polymerized at 37°C, 45°C, and 60°C for 12, 12, and 48 hours, respectively. In order to identify OHCs, semithin sections were harvested and stained with toluidine blue (Sigma-Aldrich, St. Louis, MO, USA). Ultra-thin sections of 70 nm thickness were cut in parallel with the long axis of OHCs and were collected on formvar-coated 100 mesh copper grids. Sections were contrast stained using 2% aqueous uranyl acetate (Sigma-Aldrich) and lead citrate (Sigma-Aldrich). The grids were observed, and images taken using an 80 kV transmission electron microscope (Tecnai BioTwin, FEI Compnay, Hillsboro, OR, USA) in the Electron Microscopy Facility of the Center for Cellular and Molecular Imaging (CCMI) at Yale University School of Medicine.

*TEM Tomography-.* For electron tomography, 250-nm-thick sections with 15LJnm fiducial gold to aid alignment for tomography was done using FEI Tecnai TF20 at 200LJkV. Data were collected using SerialEM (Boulder) on a FEI Eagle 4Kx4K CCD camera using tilt angles −60° to 60° (at one-degree increments) at a binned pixel size of 1.242LJnm on a 4kLJ×LJ4k Eagle CCD (FEI) using a 150 µm C2 aperture and a 100 µm objective aperture. Images were reconstructed using Imod4.9 (University of Colorado Boulder) in an automated fashion with nonlinear anisotropic diffusion filtering and manual segmentation. OHC lateral walls were reconstructed from the tomograms with ETOMO software to align stacked images by aligning gold fiducials, and segmented using IMOD software (*45–47*).

### Electromotility (eM)

OHC eM was measured in WT and αII spectrin cKO KO mice, as previously done in WT guinea pig and mouse (*48*). Mice were euthanized with an overdose of chloral hydrate (480 mg/Kg; IP), the cochleae extracted, and the organ of Corti of the apical turn dissected out in Dubecco’s Phosphate Buffered Solution (1X DPBS, with no calcium and magnesium). OHCs were then dissociated by enzymatic digestion with trypsin (0.5 mg/mL) for 5 minutes followed by gentle trituration in extracellular ionic blocking solution with a plastic pipette. Isolated cells were then transferred onto a glass bottom dish and visualized using an inverted microscope (Eclipse TI-2000, Nikon Instruments, Inc., Melville, NY, USA) with a 40X lens. Gigaohm seals were made on single isolated OHCs on the lateral membrane using 3.5-5 MΩ patch pipettes filled with intracellular ionic blocking solution. Extracellular solution contained (mM) NaCl 100, TEA-Cl 20, CsCl 20, CoCl_2_ 2, MgCl_2_ 1, CaCl_2_ 1, Hepes 10, and intracellular solution contained (mM) CsCl 140, MgCl_2_ 2, HEPES 10, and EGTA 10. The pH and osmolarity of the solutions were adjusted to 7.2 and 300 mOsm respectively. All recordings were performed at room temperature. An Axon 200B amplifier was used for whole-cell recording and an Axon Digidata 1440 was used for digitizing (both from Molecular Devices, LLC, San Jose, CA, USA). OHCs were whole-cell voltage clamped and AC analysis of membrane currents (I_m_) and eM were made by stimulating cells with a voltage ramp from 100 to -110 mV (nominal), superimposed with summed AC voltages at harmonically related frequencies of 195.3, 390.6, 781.3, 1562.5, 3125, and 6250 Hz, with a 10 μs sample clock. Currents were filtered at 10 kHz with a 4-pole Bessel filter. Corrections for series resistance and system characteristics were made during analysis. Ramp-evoked eM measures were made with fast video recording. A Phantom 310 camera (Vision Research, Inc., Wayne, NJ, USA) was used at a frame rate of 25 kHz. Magnification was set to provide 106 nm/pixel. We used our previously developed method to track the apical image (cuticular plate) of the OHC, providing sub-pixel resolution of movements (*49*). AC voltages under voltage clamp were determined by subtraction of the voltage drop across R_s_, i.e., I_Rs_ * R_s_, with I_Rs_ being the sum of resistive and capacitive components of the cell membrane current. The gains of eM (cell length change/membrane potential change; nm/mV) were based on these corrections, as we did previously (*48*).

**Supplementary Figure 1.** A Shown is the schematic for OHC specific αII spectrin cKO mice. αII spectrin fl mice were bred with prestin-CreERT2 mice to obtain homozygous αII spectrin fl/fl mice with prestin-CreERT2. Cre recombinase was then induced by Tamoxifen injection from P9-15. This results in an OHC specific deletion of exon 8 in the αII spectrin gene that is flanked by two loxP sites. B. Shows PCR products from one compound heterozygous mouse (lane 1 ) with αII spectrin fl and prestinCreERT2 that compares to a WT mouse (lane 2) and one homozygous αII spectrin fl/fl mouse with prestinCreERT2.

## Acknowledgments.

This work was funded by the NIH NIDCD R01DC016318, and R56DC021057 to JSS and DSN. We would like to thank Dr Xinran Liu, and the staff at the CCMI Electron Microscopy Core Facility, Center for Cellular and Molecular Imaging for their effort.

