## Supplementary figures and images for "Electromotility can be disassociated from gating charge movement in outer hair cells of conditional alpha2 spectrin knockout mice"

### Supplemental Figure1

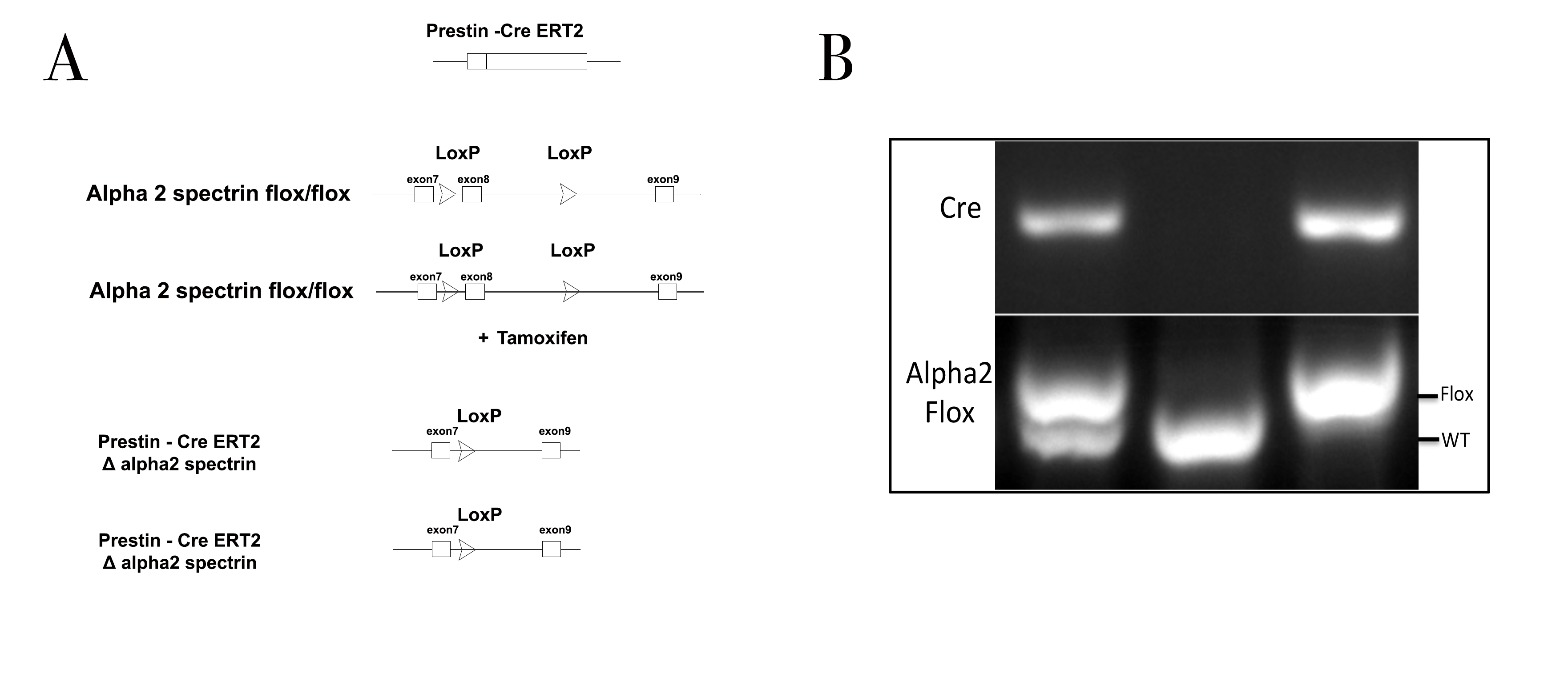
